# *Ybx1* guides C/EBPα and cBAF chromatin-remodeling complex to promote adipogenic gene expression in steatotic hepatocytes

**DOI:** 10.1101/2024.10.25.620017

**Authors:** James M. Jordan, Jixuan Qiao, Chenhui Zou, Sandra Steensels, Fahrettin Haczeyni, Alessandra Fraim, Arturro Mendoza, Ype P. de Jong, Baran A. Ersoy

## Abstract

Excessive lipid accumulation by hepatocytes underlies the pathogenesis of metabolic-dysfunction associated steatotic liver disease (MASLD) and metabolic-dysfunction associated steatohepatitis (MASH) from the earliest stages of the disease. How liver cells regulate the commitment to storing large volumes of fat despite resulting tissue damage is not well understood. Here, we show *Y box-binding protein 1* (*Ybx1*) is necessary for ectopic activation of an adipocyte-specific gene expression module that potentiates lipid accumulation in hepatocytes. Diet-induced obese (DIO) mice, with liver-specific depletion of *Ybx1* (*Ybx1*^*LKO*^), are resistant to MASLD without becoming hyperlipidemic. *Ybx1*^*LKO*^ livers exhibit upregulation of hepatocyte markers, like urea processing enzyme carbamoyl phosphate synthetase I (*Cps1)*, and downregulation of adipocyte markers known to be transcriptionally regulated by peroxisome proliferator-activated receptor gamma (*PPARγ*). In nuclei of DIO mice, YBX1 interacts with CCAAT-enhancer-binding proteins alpha (C/EBPα) and the canonical BRG1/BRM-associated factor complex (cBAF); and *C/EBPα* is required for *Ybx1*-dependent *PPARγ* expression in cultured liver cells. The chromatin binding pattern of YBX1 from DIO mouse liver overlaps with those of C/EBPα and cBAF at key adipogenic loci including *Pparg* and *Cfd*. However, most YBX1-DNA binding occurs on C/EBPα-cBAF-depleted stretches located on chromosomes 16, 18, and 19, spanning up to five Mb, and overlapping regions which are inaccessible in differentiating preadipocytes, thereby bounding activational C/EBPα-cBAF complex-DNA interactions. Moreover, YBX1 expression is increased up to nine-fold in the livers of obese patients with MASLD-MASH compared to healthy obese controls; and adipocyte-specific genes, upregulated by *Ybx1*, are also upregulated in human MASLD-MASH. Overall, our study uncovers *Ybx1* as a critical epigenetic regulator in liver and potential therapeutic target for treatment of MASLD and MASH.

## Introduction

In health, the liver periodically stores lipid to meet the demands of the body by implementing highly orchestrated transcriptional programs^1–10^. Simple steatosis represents the earliest stage of metabolic liver disease pathogenesis and is defined by the benign storage of excess lipids in droplets. Chronic lipid accumulation results in MASLD, which further progresses into MASH with irreparable liver damage^11^.

Obesity-induced hepatic lipid storage is partially driven by *PPARγ*, which plays a well-studied role as master regulator of adipogenesis in adipose tissue^12–15^. *PPARγ* activation within adipose or during the pathogenesis of simple steatosis in the liver prevents lipotoxicity through the safe storage of lipids in lipid droplets. However, hepatic *PPARγ* expression is increased in patients with MASLD^16^ and in the hepatocytes of mice with adipocyte-specific ablation of *Pparg*, which exacerbates fatty liver disease by impairing adipogenesis^17^, suggesting it contributes to MASLD pathogenesis by promoting cell autonomous lipid accumulation in hepatocytes. Therefore, the temporal and tissue-specific regulation of *PPARγ* determines its beneficial versus harmful impact.

The SWI/SNF complex, first characterized in yeast as a key chromatin-remodeling factor involved in transcriptional regulation, has mammalian counterparts known as BAF complexes^18,19^. BAF complexes are evolutionarily conserved assemblies that utilize ATP hydrolysis to modulate chromatin structure and thereby regulate gene expression^18,19^. In differentiating preadipocytes, cBAF and C/EBPα interact to promote the expression of *PPARγ*^21^. Activation of *PPARγ* is sufficient to trigger a feed-forward transcriptional loop between itself and *C/EBPα* that drives and sustains adipocyte differentiation and a commitment to an adipocyte cell fate, respectively^22,23^.

*Ybx1* is a single-stranded nucleic acid-binding protein that regulates transcription in the nucleus and mRNA processing in the cytosol with pleiotropic effects that vary with tissue and cell type^24–27^. Originally studied in its capacity as a transcriptional regulator^28–30^, it has more recently been described as a miRNA shuttle^31,32^ and m^5^C-mRNA-binding protein which post-transcriptionally regulates *Ulk1* and *Ulk2* mRNA to promote autophagy and adipogenesis^33^. It has also recently been shown to regulate thermogenesis in brown adipose tissue^34^ in its capacity as an RNA-binding protein. Moreover, *Ybx1* has been studied for its role in promoting an epithelial-to-mesenchymal transition downstream the TGFβ signaling pathway to contribute to fibrogenesis^29,35–40^.

Because genetic ablation of *Ybx1* is embryonic lethal^27^, its hepatic function in the setting of MASLD has remained incompletely understood. In this study, we generated a new mouse model with conditional ablation of *Ybx1*, to determine whether its adipogenesis-driving function within healthy fat tissue underlies its pathogenic role in the obese liver. Overall, this study shows that *Ybx1* is a powerful epigenetic regulator of hepatic metabolism and represents an attractive therapeutic target for MASLD-MASH.

## Results

### *Ybx1* is dysregulated by DIO and promotes steatosis and hyperlipidemia

Using a publicly available cohort of patient liver transcriptomes^41^, we found increased levels of *YBX1* in obese patients with MASLD-MASH (Figure 1A). Next, we measured YBX1 in liver biopsies from obese patients (BMI: 36-60) with clinically healthy liver or MASLD-MASH. YBX1 was increased by four-fold in MASLD-MASH liver (quantified in Figure 1B; immunoblot in Figure S1A). Feeding mice with high fat diet (HFD, 60% fat) for 16 w resulted in nine-fold increase in nuclear YBX1 abundance (Figure 1C). Notably, the cytosolic YBX1 fraction remained unchanged. Therefore, the pathogenic impact of *Ybx1* must originate from its function in the nucleus in the setting of MASLD.

**Figure 1:**
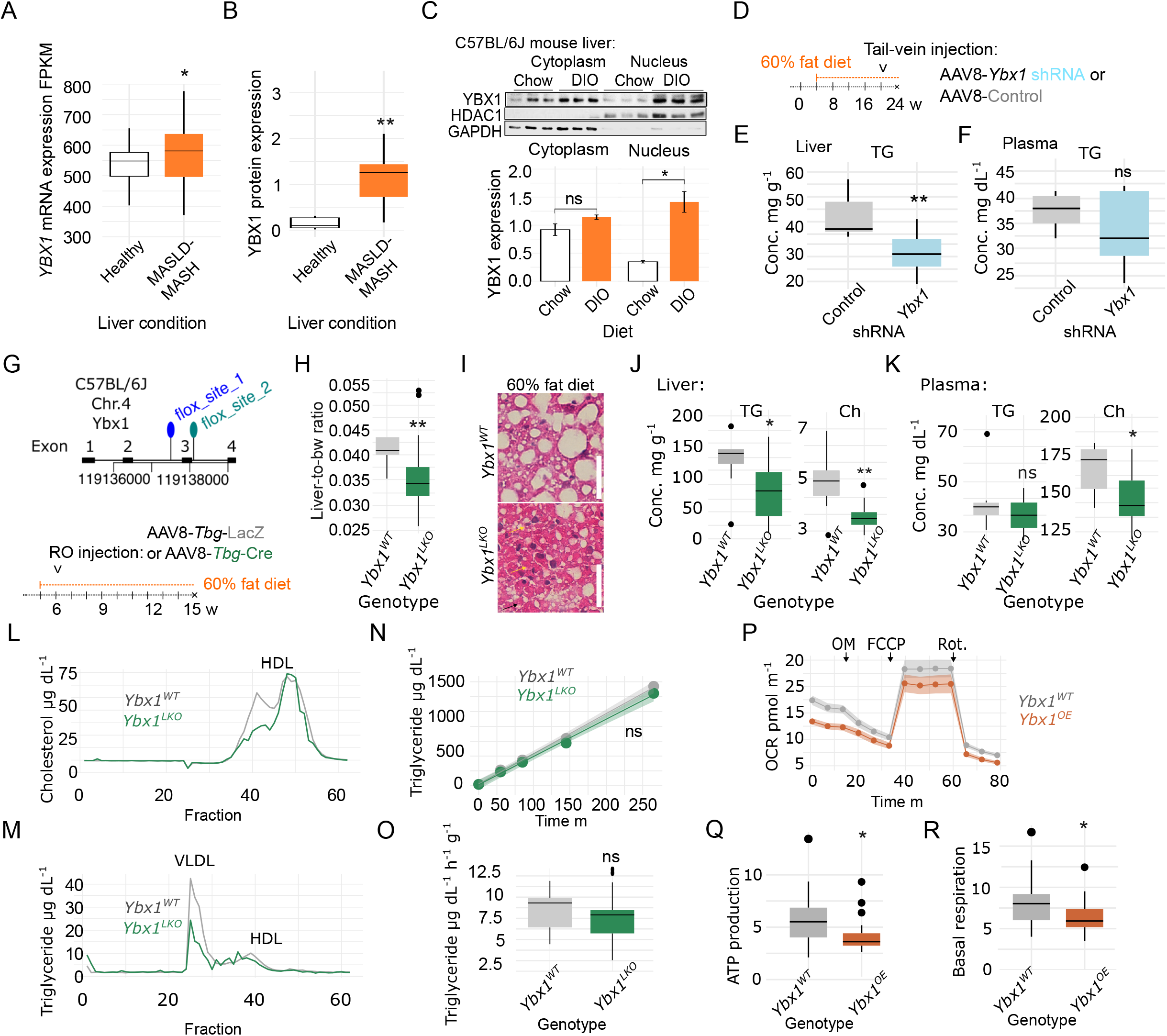
YBX1 is activated by obesity and promotes liver disease in humans and mice. A) FPKM of YBX1 in patients with MASLD-MASH compared to healthy control liver (n=26 healthy, 31 MASLD-MASH; p < 0.05, Linear Mixed Effects Model). B) Densitometric quantification of *YBX1* expression in healthy (n=6) and MASLD/MASH (n=9) livers of obese patients (36-60 BMI). C) Immunoblot and densitometric quantification of YBX1 in nuclear and cytoplasmic fractions of *YBX1* normalized to HDAC1 and GAPDH, respectively. n=3 mice/condition. D) Schematic of experimental design using *Ybx1* shRNA to reverse MASLD (vs. empty vector control) in DIO wild-type mice. E) Triglycerides in DIO (60%-fat diet) WT mouse liver and F) plasma two weeks after adenovirus-mediated *Ybx1* shRNA or empty vector control. G) Experimental design of conditional liver-specific *Ybx1* ablation in DIO mice. Exon 3 of *Ybx1* was flox’d using Crispr-*Cas9* and ablated in liver via administration of AAV8-*Tbg*-*cre* retro-orbital injection (vs. AAV8-*Tbg*-LacZ control). H) Liver-to-body weight ratio in DIO mice of given genotype. I) Representative images of H&E-stained liver sections from DIO mice. J) Biochemical analysis of whole liver and K) plasma lipids from DIO mice. L) Plasma Ch M) and TG fast-protein liquid chromatography (FPLC) measurements. N) TG secretion rate, and O) and quantification of secretion rate. n=1 pooled sample of 9 individuals/genotype. P) Oxygen consumption rate (OCR), Q) quantification of basal respiration, and R) ATP production from Seahorse mitochondrial stress test assay on mouse primary hepatocytes (mPH) of given genotypes. n=5000 cells/well, 3 wells/condition, representative of 2 trials. For B-F, H, J, K, and O, *P-value <0.05, ** < 0.01, Student’s t-test. For N, ribbon indicates +/- SEM.

To test whether the inhibition of *Ybx1* could reverse MASLD, we knocked down hepatic *Ybx1* expression in the livers of DIO mice using adenoviral delivery of shRNA targeting *Ybx1*. In mice that were fed HFD for 17 w (> 40g bw), inhibition of *Ybx1* for two weeks was sufficient to reduce concentrations of hepatic triglycerides (TG, Figure 1D) without increasing plasma TG (Figure 1F). It did not, however, affect liver cholesterol (Ch), free cholesterol (FCh), phospholipid (PL), or non-esterified fatty acid (NEFA) concentrations (Figure S1B). Plasma Ch and PL concentrations were also reduced (Figure S1C); however, we did not detect a difference in the levels of plasma FCh, or NEFA (Figure S1C).

To further these studies, we generated a mouse model with conditional deletion of *Ybx1* by inserting *LoxP* sites flanking exon three (Figure 1G). Hepatic *Ybx1* was ablated via retro-orbital injection in 5-week-old mice with adeno-associated virus (AAV8) expressing *cre* recombinase under the control of the liver-specific *thyroxin-binding globulin* (*Tbg*) promoter, which introduced a premature stop codon upon recombination. *Ybx1* was selectively ablated in the livers of DIO *Ybx1*^*fl/fl*^ mice without altering its expression within control tissues (Figure S1D). After 12 w on 60% (high)-fat diet (HFD), *Ybx1*^*LKO*^ mice did not exhibit differences in body weight (Figure S1E), nor composition (Figure S1F) compared to *Ybx1*^*WT*^ mice. However, their liver-to-body weight ratio was significantly reduced (Figure 1H), which was attributable to reduced steatosis (Figure 1I). Biochemical analysis confirmed that the liver of DIO *Ybx1*^*LKO*^ mice had markedly reduced TG, Ch, and FCh (Figures 1J, S1G). We did not detect differences in liver PL or glycogen (Figure S1G).

We next characterized plasma lipids and lipoproteins to determine whether excess TG and Ch accumulated in the plasma instead of the liver. *Ybx1*^*LKO*^ mice had reduced plasma total Ch (Figure 1K) and FCh (Figure S1H), while TG, PL, and NEFA remained unchanged (Figures 1K, S1H). Moreover, the size distribution of Ch-harboring lipoproteins was shifted by loss of hepatic *Ybx1* (Figure 1L). The LDL-rich fractions of VLDL and LDL/HDL_1_ TG were reduced in *Ybx1*^*LKO*^ while the HDL TG fraction remained unchanged (Figure 1M). Importantly, this was not due to decreased VLDL secretion from the liver (Figures 1N,O).

We hypothesized that *Ybx1* promoted the accumulation of lipids by reducing fatty acid oxidation. To test this, we measured oxygen consumption rates in primary hepatocytes. Since healthy primary hepatocytes express negligible amounts of *Ybx1*, we tested the impact of *Ybx1* overexpression on mitochondrial function. Indeed, *Ybx1* suppressed basal respiration rate and ATP production compared to controls (Figures 1P-R). Overall, these results indicate that *Ybx1* acts in the liver to promote lipid anabolism and storage at the expense of catabolic energy expenditure.

### *Ybx1* is required for a transcriptional program consistent with lipid accumulation

Out of 12,002 detected genes, we identified 371 differentially expressed genes (DEGs) between *Ybx1*^*LKO*^ and *Ybx1*^*WT*^ mice, comprising 250 downregulated and 121 upregulated genes (Figures S2A–C, File 1). Ingenuity Pathway Analysis (Qiagen) gene set enrichment analysis (GSEA) implicated disease and functions consistent with the observed hepatic phenotype (Figure 2A). DEGs were significantly enriched for Kruppel-like factor (*Klf*) and related *Sp* transcription factor occupancy sites (Figure 2B). In addition to defining cellular lineages during development, KLFs have been shown to regulate pluripotency, adipogenesis, and metabolic reprogramming^42–48^. Identifying this signature in DIO *Ybx1*^*LKO*^ liver suggests that constitutive activation of hepatic *Ybx1* affects mature hepatocyte fate in response to dietary inputs.

**Figure 2:**
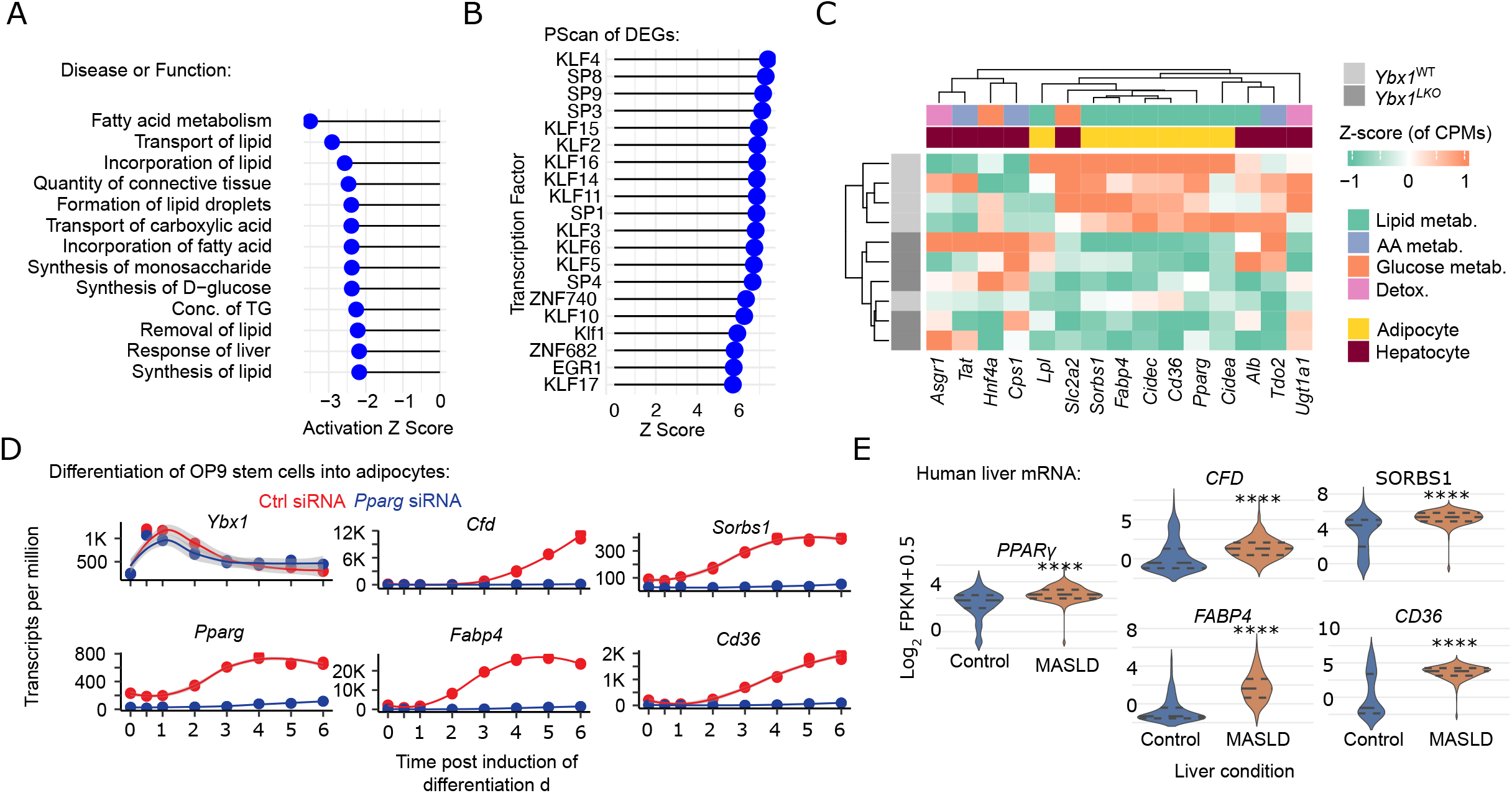
Hepatic Ybx1 activates an adipocyte-like gene expression program in the liver of DIO mice. A) Ingenuity pathway analysis (IPA; Qiagen) gene set enrichment analysis and B) promoter scan (Pscan) results from differentially expressed genes (DEGs) in *Ybx1*^*LKO*^ vs. *Ybx1*^*WT*^ whole-liver lysates derived from DIO mice. C) Heatmap of Z-score normalized counts per million reads of mRNA-seq from *Ybx1*^*LKO*^ vs. *Ybx1*^*WT*^ whole-liver lysates derived from DIO mice, of hepatocyte and adipocyte markers selected using Human Protein Atlas expression data (see Figure S2). D) Analysis of adipogenesis transcriptomic time series in *OP9* cells, with and without siRNA against *Pparg*, showing DIO *Ybx1*^*LKO*^ liver DEGs are induced in the early adipocyte differentiation. E) Violin plots of adipogenic genes expression levels in healthy (n=270) and MASLD-MASH (n=625) human liver gathered from IPA OmicsLand database.

DIO *Ybx1*^*LKO*^ mice showed reduced expression of adipocyte markers including *Pparg, Cfd, Fabp4*, and *Sorbs1*. These genes are specifically involved in adipogenesis^49–51^ and determined to be good markers via analysis of the human protein atlas database (Figure S2D). *Cd36* an acyl-CoA receptor typically restricted to adipose tissue but upregulated in steatotic liver^52–54^, was also markedly reduced in *Ybx1*^*LKO*^ liver. Conversely, hepatocyte markers *Asgr1, Tat*, and *Cps1* were depleted in *Ybx1*^*WT*^ relative to *Ybx1*^*LKO*^. Together, this suggests hepatic *Ybx1* promotes a maladaptive form of metabolic plasticity in mature hepatocytes in a setting of DIO.

Using an mRNA-seq time series of *OP9* stem cells differentiating into adipocytes^55^, we confirmed key adipogenic regulators affected by *Ybx1* in the liver are expressed *Pparg*-dependently during adipocyte differentiation (Figure 2D). Moreover, *Ybx1* expression spikes within 12 h of adipogenesis induction and peaks after 24 h, preceding the activation of *Pparg* and occurring independently of *Pparg*. To determine if these gene expression changes were relevant for human liver disease, we performed an analysis on selected genes from a transcriptomics database containing 625 MASLD-MASH liver biopsies and 270 healthy controls from the IPA OmicsLand database (Qiagen). We confirmed upregulation of key genes *PPARγ, FABP4, CFD, SORBS1*, and *CD36* (Figure 2E).

Using IPA, we confirmed traditional GSEA results by identifying three of the four known *Ppar* isoforms^56,57^ as upstream regulators (Figure 3A). Further analysis of individual and shared *Ppar* target genes indicated that *Pparg* was the predominant driver of the *Ybx1*^*LKO*^ transcriptomic signature (not shown). Genes known to be directly transcriptionally regulated primarily by *Pparg* and required for TG biosynthesis—including *Scd1, Fads2, Gpd1, Gpat3*, and *Dgat2*—were significantly downregulated (Figure 3B, Supplemental File 1). Lipid droplet proteins, and known *Pparg* targets^58^, *Cidea, Cidec*, and *Clstn3* are highly regulated by *Ybx1* in the liver. These proteins are critical regulators of lipid droplet size and dynamics, and all three are linked to MASLD^59–63^. Additional lipid droplet regulatory proteins— *Fitm1, Fitm2*, and three perilipin family members (*Plin2, Plin4*, and *Plin5*)—were also downregulated in *Ybx1*^*LKO*^ liver. Overall, our analysis suggests *Ybx1* regulates a potent adipogenic *Pparg* module in DIO hepatocytes.

**Figure 3:**
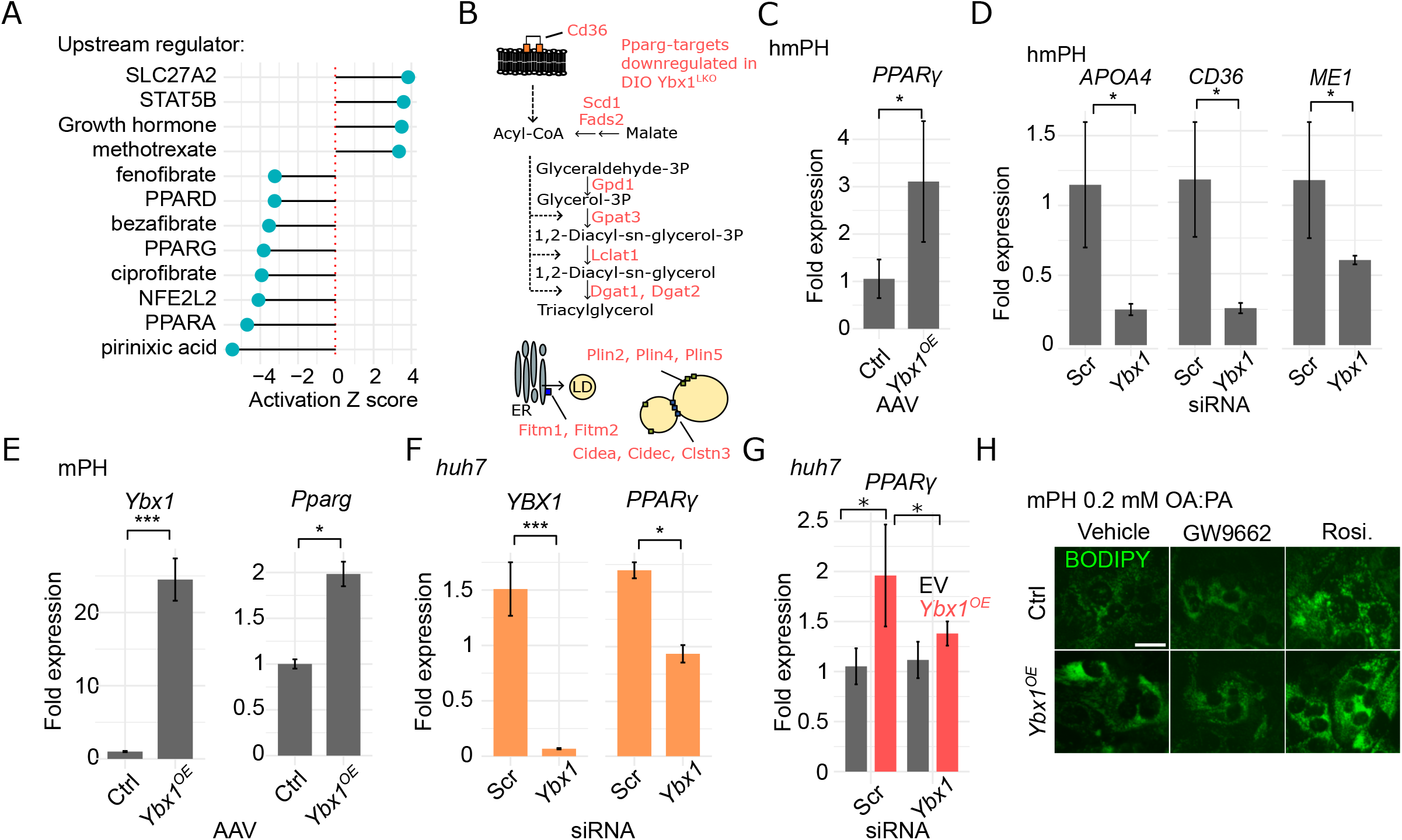
*Ybx1* is necessary and sufficient for adipogenic *Pparg* expression in hepatocytes. A) Results of IPA upstream regulator analysis using DEGs from DIO *Ybx1*^*LKO*^ and *Ybx1*^*WT*^ mouse liver. B) Depictions of *Pparg*-target genes downregulated in DIO *Ybx1*^*LKO*^ mouse liver and processes in which they are involved. C) Representative RT-qPCR results in humanized mouse hepatocytes (hmPH) with adenovirus-mediated overexpression of *Ybx1*, D) in hmPH, with siRNA knockdown of *Ybx1*, E) in mouse primary hepatocytes (mPH) with adenovirus-mediated overexpression of *Ybx1*, F) in *huh7* cells exposed to 0.5 mM bsa:palmitate (PA) for 16 h with *Ybx1* siRNA versus scrambled (Scr) siRNA control, and G) in *huh7* cells with adenovirus-mediated overexpression of *Ybx1* with and without *Ybx1* siRNA. H) Representative micrographs of BODIPY neutral lipid-stained mPH subjected to 0.2 mM bsa:oleate/bsa:PA (1:1) for 16 h with *PPARγ* antagonist GW9662 (20 μM), *PPARγ* agonist rosiglitazone (100 μM), or vehicle only. 400x magnification. For C-G, n=3 wells/condition/trial in at least 2 trials. *Huh7* and hmPH were normalized using *ActB* or *ActD*; mPH were normalized using *Cyclo*. Analyzed using the Livark method. Error bars indicate +/- SEM. *P-value<0.05, **<0.01, and ***<0.001; Student’s t-test.

Cell-autonomous adipogenic drive by *Ybx1* was confirmed in cultured cells whereby lipid droplet accumulation was suppressed by *Ybx1* siRNA treatment and increased by *Ybx1* overexpression (Figure S3A). We confirmed fatty acid treatment increased *Pparg* expression in huh7 cells (Figure S3B). In human primary hepatocytes that were harvested from mice (hmPH), *PPARγ* was increased by up to four-fold by overexpression of *Ybx1* (Figure 3C). We spot checked known *PPAR*-target genes that were differentially expressed in *Ybx1*^*LKO*^ livers. In support of similarities between *Ybx1*^*LKO*^ livers and cell culture systems, *Ybx1* siRNA suppressed *PPAR*-target genes *APOA4, CD36*, and *ME1* in hmPH (Figure 3D) and *CideC* expression in palmitate(PA)-challenged *huh7* cells (Figure S3C). Overexpression of *Ybx1* increased *Pparg* expression by two-fold in mPH (Figure 3E). In palmitate-treated *huh7* cells, *Ybx1* siRNA suppressed *Pparg* expression (Figure 3F). We confirmed enhanced *Pparg* expression was *Ybx1*-dependent by overexpressing *Ybx1* in *huh7* cells while simultaneously knocking down *Ybx1* with siRNA (Figure 3G). To link the *Ybx1, Pparg*, and hepatic lipid accumulation, we overexpressed *Ybx1* in mPH exposed to 0.2 mM bsa-conjugated oleate and PA for 16 h while treating the cells with *Pparg*-specific antagonist GW9662, *Pparg*-specific agonist rosiglitazone, or vehicle (DMSO). As expected, *Ybx1*^*OE*^ promoted lipid storage as evidenced by increased BODIPY neutral lipid staining (Figure 3H). This enhancement was negated if *Ybx1*^*OE*^ cells were treated with GW9662. Conversely, BODIPY staining was enhanced by the rosiglitazone in EV cells and further enhanced in *Ybx1*^*OE*^ cells. Overall, these results indicate that *Ybx1* is necessary and sufficient to drive the expression of *PPARγ* in liver cells, which in turn, drives an ectopic adipocyte-like gene repertoire, to promote hepatic lipid accumulation.

### YBX1 interacts with C/EBPα and cBAF in the nuclei of DIO mouse liver

To identify the molecular mechanism by which *Ybx1* affects gene expression, we immunoprecipitated (IP) endogenous YBX1 from the nuclear lysate of mouse liver with MASLD and performed mass spectrometry-based proteomics to identify YBX1-binding proteins in the context of MASLD. Analysis suggests YBX1 is closely associated with several varieties histone H2, as well as C/EBPα, and six members of the BAF nucleosome remodeling complex including canonical BAF (cBAF)-specific DPF2^64^ (Figure 4A, Figures S4A,B). Of the 19 most significantly enriched YBX1 coIP hits, only *Cebpa* and *Cebpb* are known to directly transcriptionally regulate *Pparg* expression (Figure S4C). In differentiating pre-adipocytes, *cBAF* and *C/EBPα* promote the transcription of *PPARγ*^22,23,65–70^; and *C/EBPα* expression is increased in MASLD^71,72^. We found that in the absence of *C/EBPα, PPARγ* was not induced in response to PA exposure (Figure S4C). Combining *YBX1* and *C/EBPα* siRNA did not result in an additive effect on the negative regulation of *PPARγ* expression (Figure S4D). Next, we overexpressed *Ybx1* in mouse primary hepatocytes while knocking down *Cebpa* with siRNA. We found that *Ybx1*-dependent *Pparγ* expression required *Cebpa* (Figure 4B). Finally, the knockdown of *Ybx1* and *Cebpa* reduced neutral lipid staining in mouse primary hepatocytes to similar extents (Figure S4E). Taken together, these results demonstrate that *C/EBPα* requires *Ybx1* to induce *PPARγ* expression and lipid accumulation in DIO hepatocytes.

**Figure 4:**
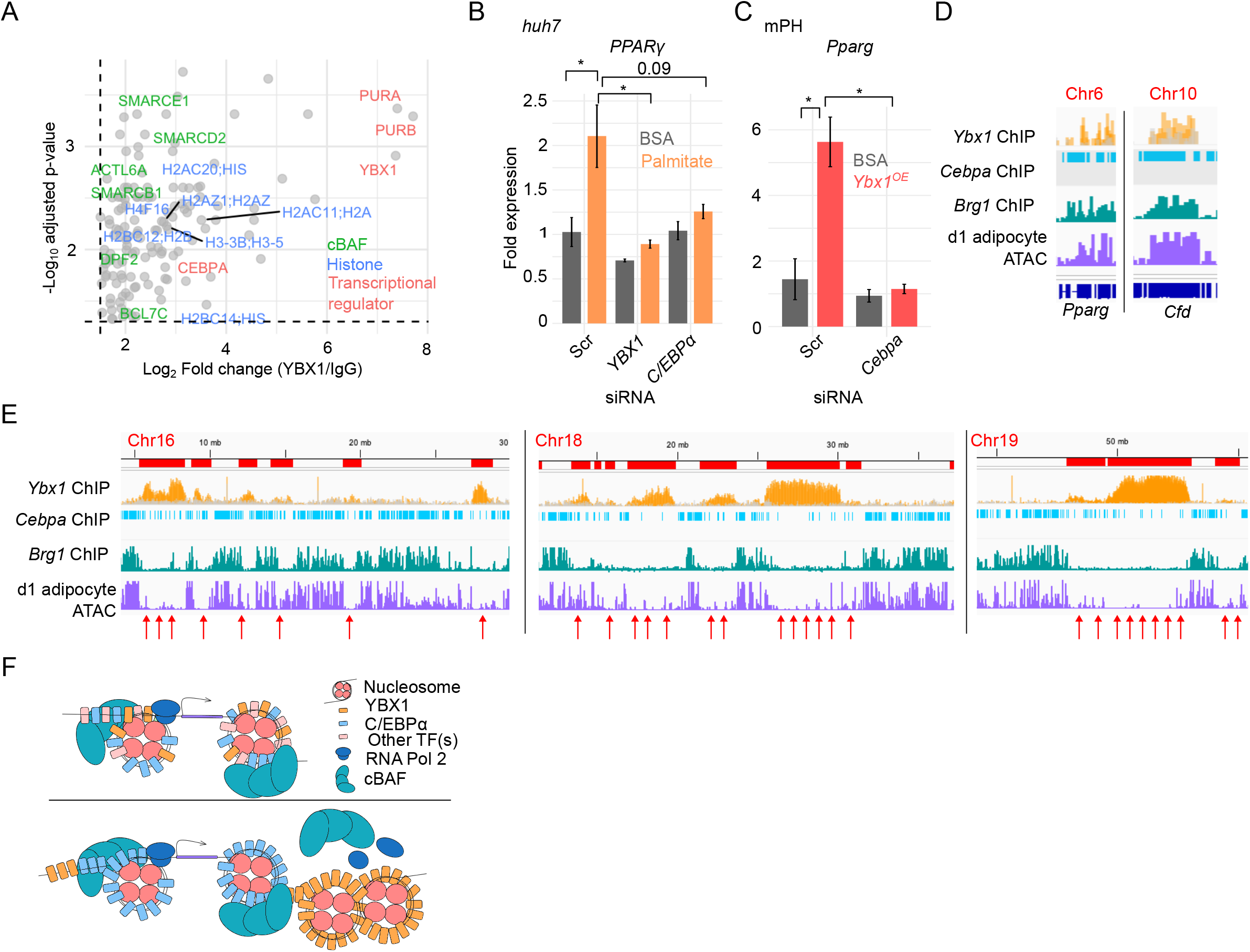
Nuclear YBX1 regulates C/EBPα-cBAF-DNA interactions to shape the chromatin landscape in a setting of diet-induced obesity. A) Volcano plot showing histones, cBAF complex members, and transcriptional regulators enriched with coIP of YBX1 from the nuclei of DIO liver and quantified with LC-MS/MS. B) Representative RT-qPCR results in *huh7* cells exposed to 0.5 mM bsa:PA or bsa-only control for 16 h with siRNA against *YBX1, C/EBPα*, or scrambled (Scr) siRNA control, and C) in *huh7* cells exposed to 0.5 mM bsa:PA or bsa-only control, with *Cebpa* and Scr siRNA. D) IGV tracks of chIP-seq analysis of liver from DIO *Ybx1*^*WT*^ mice over input (yellow) and DIO *Ybx1*^*KO*^ over input (gray); *Cebpa* chIP-seq profile from mouse liver (GSM427088), *Brg1* (cBAF subunit) chIP-seq in differentiating mouse cells (GSM7770720) and *3t3* mouse epithelial cell ATAC-seq on day 1 of induced adipogenesis (GSM5174367) at the *Pparg* locus on chr 6 and *Cfd* locus on chr 10, and for E) relatively large sections of chr 16, 18, and 19 in C/EBPα-cBAF-depleted regions corresponding to regions inaccessible in differentiating preadipocytes (red arrows). F) Working model of two distinct modes by which YBX1 regulates C/EBPα-cBAF chromatin interactions. (top) YBX1, C/EBPα, and (an) other transcription factor(s) recruit cBAF to specific nucleosomes. (bottom) Without the involvement of other TFs that allow for co-binding of YBX1 and C/EBPα-cBAF, or perhaps when other combinations of TFs are involved, YBX1-DNA interactions restrict where activational C/EBPα-cBAF interactions can occur – effectively bounding accessible chromatin in differentiating cells. For B and C, n=3 wells/condition/trial in at least 2 trials. *Huh7* were normalized using *ActB*. Analyzed using the Livark method. Error bars indicate +/- SEM. *P-value<0.05, **<0.01, and ***<0.001; Student’s t-test.

### YBX1, C/EBPα, and cBAF shape the chromatin landscape in DIO hepatocytes

To determine the molecular mechanism by which YBX1, C/EBPα, and cBAF regulate adipogenic gene expression in hepatocytes, we performed chromatin immunoprecipitation and sequencing (ChIPseq) of YBX1 in DIO *Ybx1*^*WT*^ and *Ybx1*^*LKO*^ liver, and compared the resulting binding patterns to *Cebpa* ChIPseq profiles acquired from mouse liver^73^, *cBAF* (*Brg1* subunit) ChIPseq from differentiating mouse cells^74^, and accessible chromatin regions in mouse preadipocytes on the first day of adipocyte differentiation^75^. As expected, the genomic profiles of bound cBAF and C/EBPα were highly correlated with accessible chromatin regions in differentiating preadipocytes – indicative of an important role shaping the adipogenic chromatin landscape (Figure S4). Intriguingly, the ChIP profile of *Ybx1* in DIO mouse hepatocytes suggests two distinct modes by which YBX1 interacts with cBAF and C/EBPα to affect adipogenic chromatin accessibility. Overall, YBX1 is sparsely bound to hepatic DNA; however, we identified co-binding of all three factors to adipogenic master regulators *Pparg* and *Cfd* - which are also accessible in d1 preadipocytes (Figure 4D). Most *Ybx1* ChIP reads mapped to inter- and intragenic regions spanning up to 5 Mb that were predominantly on chromosomes 16, 18, and 19 (Figure 4E). These regions are uniformly flanked by but otherwise devoid of cBAF and C/EBPα. These findings indicate YBX1 can bind at specific loci with cBAF and C/EBPα to promote accessibility, but in most cases, shapes the chromatin landscape by excluding C/EBPα-cBAF from relatively long regions of chromatin (modeled in Figure 4F).

## Discussion

In this study, we identified hepatic *Ybx1* as a critical regulator of liver adipogenicity. DIO mice lacking hepatic *Ybx1* are resistant to hepatic steatosis and other indicators of metabolic disease. We show obesity-induced activation of *Ybx1* within hepatocytes transcriptionally reprograms cells via C/EBPα-cBAF complex interactions occurring at high-level regulators of adipocyte differentiation. Moreover, *Ybx1* appears to reshape the chromatin landscape of hepatocytes by defining active cBAF-C/EBPα boundaries which effectively mimics the epigenetic landscape of differentiating preadipocytes. Reorganization of chromatin and enhancement of *PPARγ* and *Cfd* expression is sufficient to explain the striking impact on metabolic phenotypes in DIO *Ybx1*^*LKO*^ mice. In addition to directly promoting other adipocyte differentiation-related genes, *PPARγ* directly regulates metabolic gene expression to shunt away from mitochondrial oxidation and toward TG biosynthesis. *PPARγ* also directly promotes the transcription of *CD36* and lipid droplet regulatory proteins to facilitate lipid import and droplet enlargement, respectively. Interestingly, we find this enhanced adipogenic gene expression negatively correlates with hepatocyte-specific gene expression such as that of urea processing enzyme *Cps1*. Thus, *Ybx1* acts as a thrifty gene^76^ - activating a maladaptive nutrient storage program in hepatocytes at the expense of healthy hepatocyte function such as urea processing.

*C/EBPα* and *cBAF* have been shown to regulate chromatin accessibility in other tissues^77,78^ including at the *PPARγ* locus in differentiating preadipocytes^21^. *Ybx1* expression spikes five-fold shortly after exogenous induction of adipocyte differentiation independent of and preceding *PPARγ* expression (Figure 2F using data from^55^). Consistent with a role in ectopic activation of adipogenesis in mature hepatocytes, Y*bx1* has recently been implicated in regulating adipogenesis^33,34^. Comparative analyses between hepatic steatosis and adipogenesis in the presence and absence of *Ybx1* will help us understand the extent to which “trans-differentiation” of hepatocytes explains the stages of MASLD pathogenesis.

*PPARγ*-driven impact of *Ybx1* was the primary focus of this study due to reduction of nearly all *PPARγ* targets as well as *PPARγ* itself. Loss of hepatic *Ybx1* did not prevent all metabolic dysregulation-associated lipogenic events as other HFD-associated pathways, especially sterol response-element binding protein 1 (*Srebf1*), were enhanced in *Ybx1*^*LKO*^ relative to *Ybx1*^*WT*^ (Supplementary File 1). While this points to *PPARγ*-specific regulation, the loss of *Ybx1* also altered *PPARα* activity, which has overlapping targets with *PPARγ*, and *Pparγ* positively regulates *Pparα* expression in DIO mice^79^. *Ybx1* altered *PPARα*-specific targets in ways that were inconsistent with deactivation of *PPARα* whereas targets positively regulated solely by *PPARγ* or upregulated by *PPARα* and/or *PPARγ* were uniformly reduced in the absence of hepatic *Ybx1* (Figure S3). Nonetheless, it cannot be ignored that there is a strong signature of *PPARα* activity being altered by loss of *Ybx1*, and some genes affected by *Ybx1* related to cholesterol transporters and bile acid-related cytochromes are known to be direct targets of *PPARα* and not *PPARγ* ^80^. Finally, *Ybx1* is also known to bind and inhibit RNA in cancer or continuous cell lines^32,81,82^. This function, however, has been linked to cytosolic sequestering of mRNA by cytosolic YBX1 while our observations in HFD-fed mice placed YBX1 within the nucleus. Therefore, we conclude that obesity-activated hepatic *Ybx1* is primarily a *cBAF-C/EBPα* regulator that mediates chromatin modifications to produce maladaptive *PPARγ*-dependent lipid accumulation. These findings provide important insight into environmental cellular reprogramming of mature liver cells and suggest *Ybx1* is a promising therapeutic target for the treatment of metabolic diseases.

## Author contributions

JMJ and JQ conducted most of the experiments and analyzed the data. SS, FH, and BAE conducted experiments. JQ and AM conducted the ChIP-seq experiment and JMJ analyzed the data. CZ and YdJ contributed humanized mouse primary hepatocytes to the project. JMJ and BAE wrote the manuscript. BAE conceived and supervised the project.

## Acknowledgments

We would like to thank Dr. David Cohen, MD, PhD (Brigham and Women’s Hospital, Boston, MA) and his lab members for helpful discussions about this work. We would also like to thank Kathleen Corey, MD (Fatty Liver Clinic, Massachusetts General Hospital) for providing us with human liver biopsy samples; Guoan Zhang, PhD, and the WCM Metabolomics Core facility (New York, NY), for conducting LC-MS/MS; and the core services of the Laboratory of Comparative Pathology at the Center of Comparative Medicine & Pathology (Memorial Sloan Kettering Cancer Center, New York, NY).

## Methods

### Mice

Mouse protocols were approved by the Institutional Animal Care and Use Committee of Weill Cornell Medical College. Mice were housed under a 12-hour light/dark cycle in a climate-controlled environment. *Ybx1* conditional liver knock-out model (*Ybx1*^*LKO*^)- LoxP sites were inserted into the introns flanking *Ybx1* exon 3 using CRISPR-*Cas9* in WT C57BI/6J as previously described 5 w old *Ybx1*^*flox/flox*^ mice were injected retro-orbitally with adeno-associated virus vector serotype 8 (AAV8) expressing *cre recombinase* under transcriptional control of the hepatocyte-specific thyroxine-binding globulin (*Tbg*) promoter (*Ybx1*^*LKO*^; 5×10^10^ GC/mouse, Vigene Biosciences, Rockville, MD)^83^. Control mice were injected with AAV8-*Tbg*-*LacZ* (*Ybx1*^*WT*^, 5×10^10^ GC/mouse, Vigene Biosciences). Mice were fed a HFD (60% kcal from fat; Research Diets, D12492, New Brunswick, NJ) starting at 4-w old until they were sacrificed at 16 w old.

*Ybx1* shRNA was performed in 4w-old, wild-type (C57BI/6J) mice, fed HFD for 17 w prior to administration of shRNA and then after shRNA as well. Mice were injected retro-orbitally with AAV8-*Ybx1* shRNA or empty vector control. Physiological experiments were conducted between 7-10 d post injection and mice were sacrificed 2 w post injection.

Mice were fasted for 6 h during the light cycle before blood and tissue collections. Animals were sacrificed by exsanguination or cervical dislocation after being anesthetized with isoflurane or ketamine according to standard techniques. Blood was collected via cardiac puncture and plasma was separated by centrifuging samples at 3,000 rpm for 15 min at 4°C with ethylenediaminetetraacetic acid (EDTA). Tissue samples were snap-frozen in liquid nitrogen, immediately after collection. All samples were stored at −80 °C.

### Body composition

Fat mass and lean mass were measured in live, non-anesthetized mice using EchoMRI 3-in-1 Body Composition Analyzer as previously described^84^ (EchoMRI LLC, Houston, TX). Briefly, fasted mice were transferred into an animal holding tube and immobilized gently with a plunger to complete a 100-second scanning procedure.

### VLDL secretion rates

Mice were fasted for 5 h during the light cycle before retro-orbital injection with the lipoprotein lipase inhibitor Tyloxapol (500 mg/kg body weight) (MilliporeSigma) (3). Tail blood samples were collected at 0.5, 1, 2, and 4 h after injection. Samples were mixed with EDTA before plasma collection. Plasma TG concentrations were measured using a colorimetric enzymatic assay kit according to the manufacturer’s instructions (Wako Diagnostics, Mountain View, CA). Secretion rates were calculated as previously described^84^.

### Liver histology

Liver samples were frozen in plastic cryomolds with O.C.T. compound (Sakura Finetek, Torrance, CA) on dry ice. Samples were then fixed, sectioned, and stained with hematoxylin and eosin (H&E) or F4/80 using standard techniques. Images were captured using an Eclipse Ti microscope (Nikon, Melville, NY).

### Hepatic lipid extraction

50 mg frozen liver was homogenized in 1.5 mL methanol in a 3 mL Potter-Elvehjem homogenizer using a PTFE pestle. Lipids were extracted with 7.5 mL of chloroform/methanol (2:1, v/v). 1.5 mL 0.05% H_2_SO_4_ (1:5, v/v) was added for phase separation and the mixture was incubated at room temperature overnight. 1 mL of the bottom phase was mixed with 0.5 mL 3% triton X-100 (MilliporeSigma), dried under nitrogen gas, and resuspended in 0.5 mL water before downstream analysis.

### Hepatic glycogen

10 mg frozen liver tissue was minced and then hydrolyzed in 250 μL 2M HCl at 100°C for 1 h with periodic vortexing in 10 m intervals. 250 μL 2M NaOH was added to neutralize the hydrolysis and then centrifuged at 22,000 x g for 10 m.

### Plasma lipids

The concentrations of plasma TG, NEFA, total Ch, free Ch, and PL were measured using enzymatic kits according to the manufacturer’s instructions (Wako Diagnostics, Mountain View, CA).

### Plasma lipoproteins

Plasma was fractionated by fast protein liquid chromatography (FPLC) using a Superose 6 HR10/30 column (Pharmacia Biotech, Piscataway, NJ). 250 μL of pooled plasma was loaded into a PBS equilibrated column and eluted into 0.3 mL fractions at a flow rate of 0.3 mL/min. Fraction cholesterol and TG concentrations were measured using enzymatic kits as described above. Plasma concentrations were calculated as products of plasma TG or total Ch concentrations within relative FPLC peak areas of the corresponding lipoprotein fraction as described previously^84^.

### Immunoblot analyses

Cells were washed with ice-cold PBS and lysed in RIPA buffer with protease and phosphatase inhibitors (MilliporeSigma) for 15 m with agitation at 4°C. Mouse tissues were homogenized in RIPA buffer using a Bead Ruptor 24 Elite bead mill homogenizer (Omni International, Kennesaw, GA). Samples were then centrifuged at 16,000 x g for 20 min at 4°C. Protein concentrations were determined by Pierce BCA protein assay kit according to the manufacturer’s protocol (ThermoFisher). Protein samples were denatured in Laemmli buffer at 96°C for 5 min and then equivalent amounts were loaded into SDS-PAGE gels for separation and transferred to nitrocellulose membranes (Cytiva, Marlborough, MA) using semi-dry transfer (BioRad, Hercules, CA) using standard techniques. Membranes were blocked in TBS-T buffer (20mM Tris, 0.05% Tween-20 (v/v), pH 7.4) with 5% non-fat dry milk (w/v) and 1% bovine serum albumin (BSA, w/v, ThermoFisher) for 1 h at room temperature and then incubated with primary antibody at 4°C overnight. Membranes were washed three times with TBS-T for 5 min and incubated for 1 h at room temperature with anti-rabbit immunoglobulin/HRP secondary antibody (Agilent, Santa Clara, CA) or with anti-mouse IgG-Peroxidase secondary antibody (MilliporeSigma), followed by three washes with TBS-T (5 min each). The western blot was detected by chemiluminescent substrates (ThermoFisher) and imaged using ChemiDoc Imaging systems (BioRad, Hercules, CA). Relative protein expression was quantified by densitometry analysis using ImageJ.

### mRNA sequencing library preparation

Libraries were prepared from frozen total RNA samples from five HFD-fed *Ybx1*^*WT*^ and *Ybx1*^*LKO*^ mice. In brief, mRNA was purified using poly-T oligo-attached magnetic beads, fragmented, and then cDNA was synthesized using random hexamer primers. Next, second-strand cDNA synthesis was performed followed by end repair, A-tailing, adapter ligation, size selection, amplification, and purification. Library quality was checked with Qubit, quantified with real-time PCR, and bioanalyzed to detect size distribution. Libraries were sequenced on Illumina NovaSeq 6000 by Novogene (San Diego, CA).

### Differential gene expression analysis

Differential gene expression analysis was conducted using R. Paired-end reads were aligned using rhisat2 onto the ‘BSgenome.Mmusculus.UCSC.mm10’ with the qAlign function in the QuasR package^85^. Gene levels were counted using the qCount function in QuasR. Differential gene expression analysis was performed on protein-coding genes with at least one count per million in a sample (n=12002 detected) using an exact test in edgeR^86^. Genes with a false discovery rate (FDR) of less than 0.05 and a log_2_ fold change < −0.5 or > 0.5 were considered differentially expressed.

### Enrichment analyses

For Ingenuity Pathway Analysis (IPA; Qiagen), we applied a filter of FDR < 0.05 and log_2_ fold change < −0.5 or > 0.5. This filtering resulted in 364 genes (245 down and 119 up), subsequently used in IPA canonical pathway and upstream regulator analysis tools. Pscan was implemented on this set of DEGs as described by its authors^87^.

### RNAseq heatmaps

Heatmaps were generated using the R package ComplexHeatmaps^88^. CPMs were scaled to generate Z scores, genes were subset based on analysis of expression data from The Human Protein atlas and KEGG pathways, and finally, unsupervised hierarchical clustering was performed.

### Adenovirus production

*Ybx1* overexpression (*Ybx1*^*OE*^) and shRNA knockdown adenoviruses were generated using the Ad Easy Adenoviral Vector System (Agilent, Santa Clara, CA). *Ybx1* cDNA was amplified from MGC Mouse *Ybx1* cDNA plasmid (GE Healthcare Dharmacon, Chicago, IL) by PCR (see Table S1 for primers) and then cloned into pShuttle-CMV vector. *Ybx1* shRNA was amplified from MISSION shRNA plasmid (MilliporeSigma) and then cloned into pShuttle-CMV vector (Addgene). Plasmids containing *Ybx1* cDNA or shRNA, and empty vector were linearized with the restriction enzyme PmeI (New England Biolabs, Ipswich, MA) and transformed into BJ5183 *Escherichia coli* (Agilent, Santa Clara, CA) to produce recombinant adenovirus plasmid. Human embryonic kidney 293AD (HEK293AD) cells were maintained in DMEM with 10% fetal bovine serum (FBS), and 1% penicillin-streptomycin (PS) media (10,000 U/mL ThermoFisher). 1 μg of PacI (New England Biolabs, Ipswich, MA) linearized recombinant adenovirus plasmid was transfected to HEK293AD cells using Lipofectamine 2000 (ThermoFisher) at 70% confluency in a 6-well plate to prepare primary adenovirus stock. Adenovirus amplification was achieved by several infections of HEK 293AD cells. Cell suspension was subjected to four rounds of freeze/thaw between −80°C freezer (10 m) and 37°C water bath (10 min) and then incubated with benzonase nuclease (MilliporeSigma) for 30 min at 37°C to release viral particles. Cell lysate was collected by centrifugation at 3,200 x g for 20 min at 4°C. Particles were then purified using a cesium chloride (CsCl) gradient. Briefly, 7 mL 1.25 g/mL CsCl, 7 mL 1.4 g/mL CsCl and 22 mL cell lysate were layered from bottom to top in 38.5 mL tubes (Beckman Coulter, Brea, CA) and centrifuged at 110,880 x g for 1 h at 4°C. The bottom band was collected and diluted with the same volume of 50 mM Tris (pH 7.8), 10 mM MgCl_2_. 24 mL CsCl and the diluted viral suspension were layered from bottom to top in 38.5 mL tubes and centrifuged at 75,000 x g for 20 h at 4°C. Viral suspension was collected and dialyzed into 10 mM Tris (pH 7.8), 10 mM MgCl_2_, 88 mM sucrose and 150 mM NaCl. *Ybx1*^*OE*^ particles were purified using Adenovirus Purification Kit (Virapur, San Diego, CA) according to the manufacturer’s instructions. Titer was measured using the Adeno-X Rapid Titer Kit (Takara Bio, Mountain View, CA) according to the manufacturer’s instructions. The virus was stored at −80°C.

### Cell culture

Cells were cultured using standard techniques. *Huh7* cells were cultured in Dulbecco’s Modified Minimal Medium (DMEM, ThermoFisher) with 10% fetal bovine serum (FBS, Gennesee Scientific) and 1% penicillin-streptomycin (PS, ThermoFisher). Mouse primary hepatocytes were isolated using standard techniques as previously described^84^ and were cultured in William’s E medium (ThermoFisher) with 10% FBS and 1% PS. Humanized mouse primary were produced and harvested as previously described and then cultured in Hepatocyte Culture Medium (HCM) (Lonza Biosciences). All cells were incubated at 37°C and 5% CO_2_.

### siRNA

siRNA experiments were conducted using Lipofectamine RNAiMAX transfection reagent (ThermoFisher) according to the manufacturer’s instructions. *Huh7* cells were forward transfected, and primary hepatocytes were reverse transfected. Experiments were conducted within 72 h of transfection. See key resource table for specific siRNA used in this study.

### Fatty acid supplementation

Palmitic acid (MilliporeSigma) and oleic acid (Cayman Chemicals) were conjugated onto fatty acid-free bovine serum albumin (BSA, MilliporeSigma). 5-10 mM stocks were diluted in cell medium and added to cells. *Huh7* cells were exposed to 0.5 mM PA for 16 h. mPH and humanized mPH were exposed to 0.5 mM PA for 6 h or to 0.2 mM OA:PA (1:1) for 16 h.

### Neutral lipid staining

10 nM 500/510 BODIPY (ThermoFisher) was co-administered with fatty acid treatment. Cells were washed twice in phosphate buffered saline (PBS), fixed in 4% paraformaldehyde for 30 m at room temperature, washed twice more with PBS, and then imaged at 400x magnification with a fixed exposure time.

### RT-qPCR

Cells were lysed in Trizol (ThermoFisher) for 5 minutes and then RNA was purified using Direct-zol RNA Miniprep kit according to the manufacturer’s instructions (Zymo). RNA was reverse transcribed using UltraScript cDNA synthesis kit (PCR Biosystems). RT-qPCR was performed using Power SYBR green master mix according to the manufacturer’s instructions (ThermoFisher) with 5μL of cDNA (1:10 diluted) per reaction on a QuantStudio6 (ThermoFisher) instrument. See key resource table for qPCR primers used in this study.

### Hepatic nuclei isolation and protein extraction

Freshly harvested livers were dounced on ice until homogenous in 10 mL 250 mM STM buffer. Cell pellets were washed twice by centrifuging at 800 x g for 15 m at 4°C in 250 mM STM buffer. Pellets were resuspended in 9 mL of 2M STM buffer and then filtered through several layers of cheesecloth before being gently pipetted onto 4 mL 2M STM in ultracentrifuge tubes. Samples were centrifuged at 80000 x g for 35 m at 4°C. Nuclear pellets were then washed in cold PBS and resuspended in nuclear extraction buffer. After rocking for 20 minutes at 4°C, nuclei were lysed with 10 passes through an 18-gauge needle and sonication (30% power for 20 s, twice). Finally, samples were centrifuged at 16000 x g at 4°C for 20 m.

### Coimmunoprecipitation of YBX1 for proteomics

Nuclear protein lysates were adjusted to 1 μg/mL in cold RIPA buffer with protease and phosphatase inhibitor tablets (Invitrogen). Samples (1 mL each) were then incubated overnight at 4°C with 2 μg of primary antibody (see key resource table for antibodies). Next, samples were allowed to reach room temperature, and then 100 ul of Dynabeads (Invitrogen) were added to each sample. Samples were rotated for 1 h at room temperature, washed according to manufacturer’s instructions, and eluted in non-denaturing buffer at 37°C with rotation for 5 m.

### Tandem mass spectrometry

30 μl of coIP eluates were briefly run on a 6%/15% acrylamide gel to separate most protein from the intact antibody. Gels were stained in Coomassie blue using standard techniques, and then gel slices containing IP’d protein were excised. Protein was digested with trypsin in gel, stage-tip desalted, and then LC-MS/MS was performed. Samples were analyzed using a data-independent acquisition method and identified using the UniProt mouse protein database.

### *Ybx1* chromatin immunoprecipitation and sequencing (ChIP-seq)

ChIP-seq was performed to assess YBX1 chromatin binding in four DIO *Ybx1*^*WT*^ mice and four DIO *YBX1*^*LKO*^ animals to control for false-positive binding events. Input chromatin samples were collected from each animal prior to immunoprecipitation for use as a control. Briefly, liver from DIO mice were isolated and cross-linked in 1% formaldehyde for 10 m at room temperature to preserve protein-DNA interactions. The cross-linking reaction was quenched with 0.125 M glycine for 5 m. Chromatin was then isolated by lysis and sonicated to an average size of 200– 500 bp using a Bioruptor (Diagenode). For each sample, chromatin was IP’d overnight at 4°C with 2 μg of anti-YBX1 antibody (Abcam, ab12148) pre-bound to Dynabeads Protein G (ThermoFisher). IP’d chromatin was then washed sequentially with low-salt, high-salt, and LiCl buffers, followed by TE buffer washes. DNA-protein complexes were eluted, and cross-linking was reversed by incubating samples at 65°C overnight. The resulting DNA was purified using the QIAquick PCR purification kit (Qiagen). Library preparation was performed using the NEBNext Ultra II DNA Library Prep Kit (New England Biolabs) according to the manufacturer’s instructions. Sequencing was performed on an Illumina NovaSeq 6000 platform (by Novogene), yielding paired-end 150 bp reads.

### ChIP-seq analysis

Analysis was performed on Y*bx1*^*LKO*^ and *Ybx1*^*WT*^ mouse samples with their respective input controls. Raw paired-end sequencing reads were aligned to the *mm10* mouse reference genome using Bowtie2. Aligned reads were sorted, indexed, and coverage tracks were generated with deepTools for visualization. Additional tracks used in analysis were acquired from GEO (day 1 3t3 ATAC: GSM5174367; liver Cebpa ChIP-seq: GSM427088; and Brg1 in differentiating mouse cells ChIP-seq: GSM7770720). Tracks were visualized using IGV_2.18.2 using the GRCm38/mm10 genome as a reference.

---

**Figure S1.**
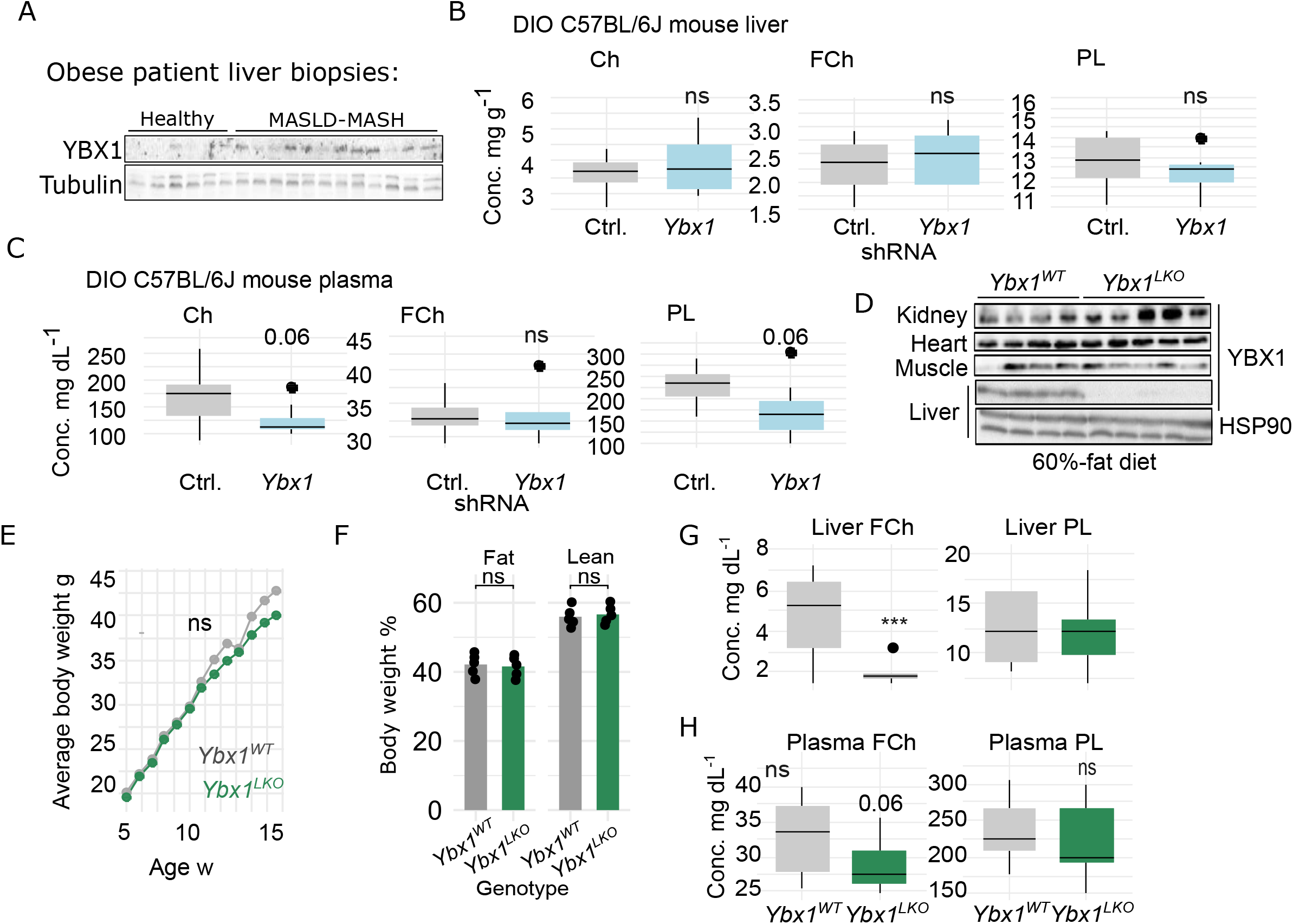

**Figure S2.**
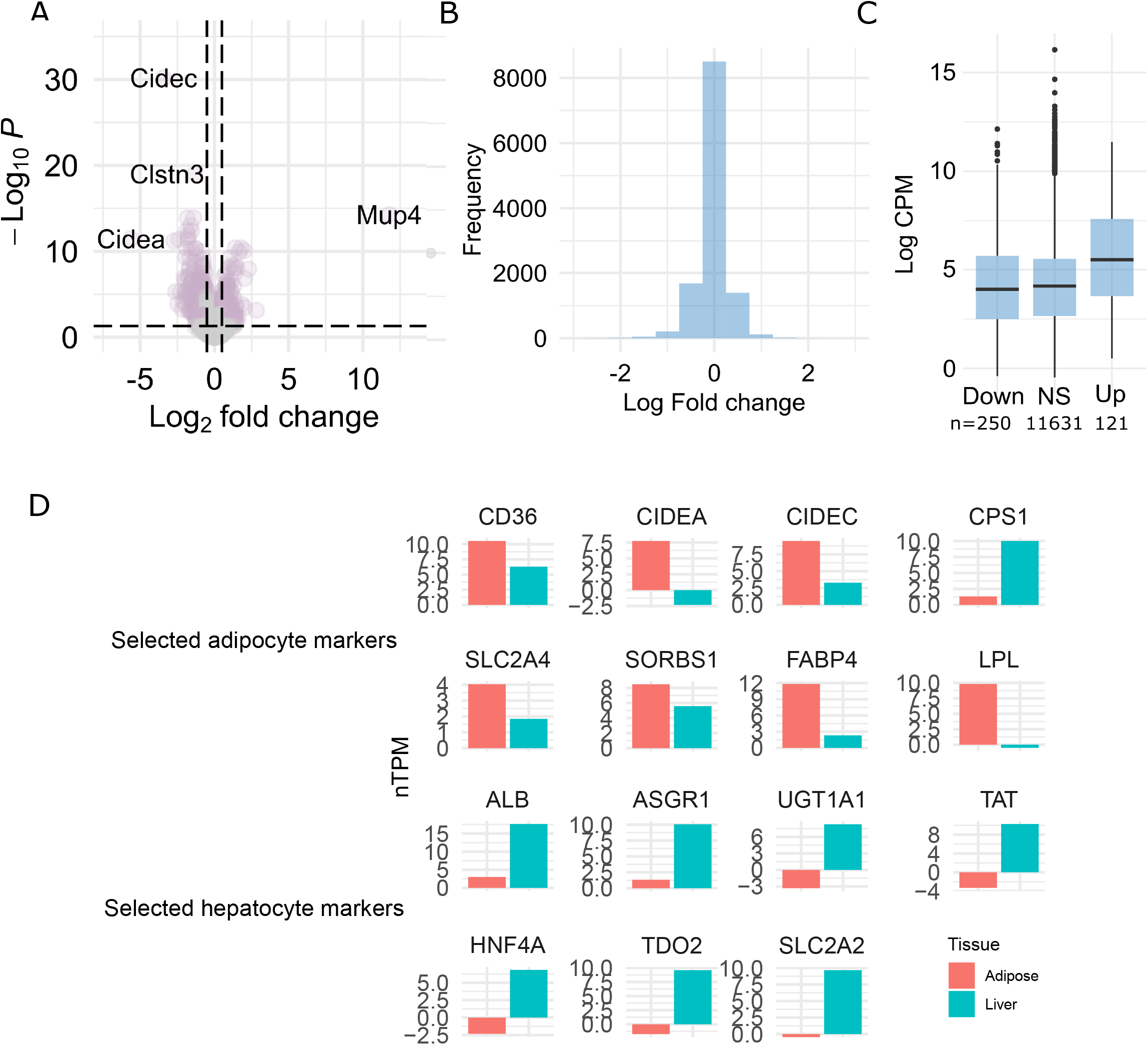

**Figure S3.**
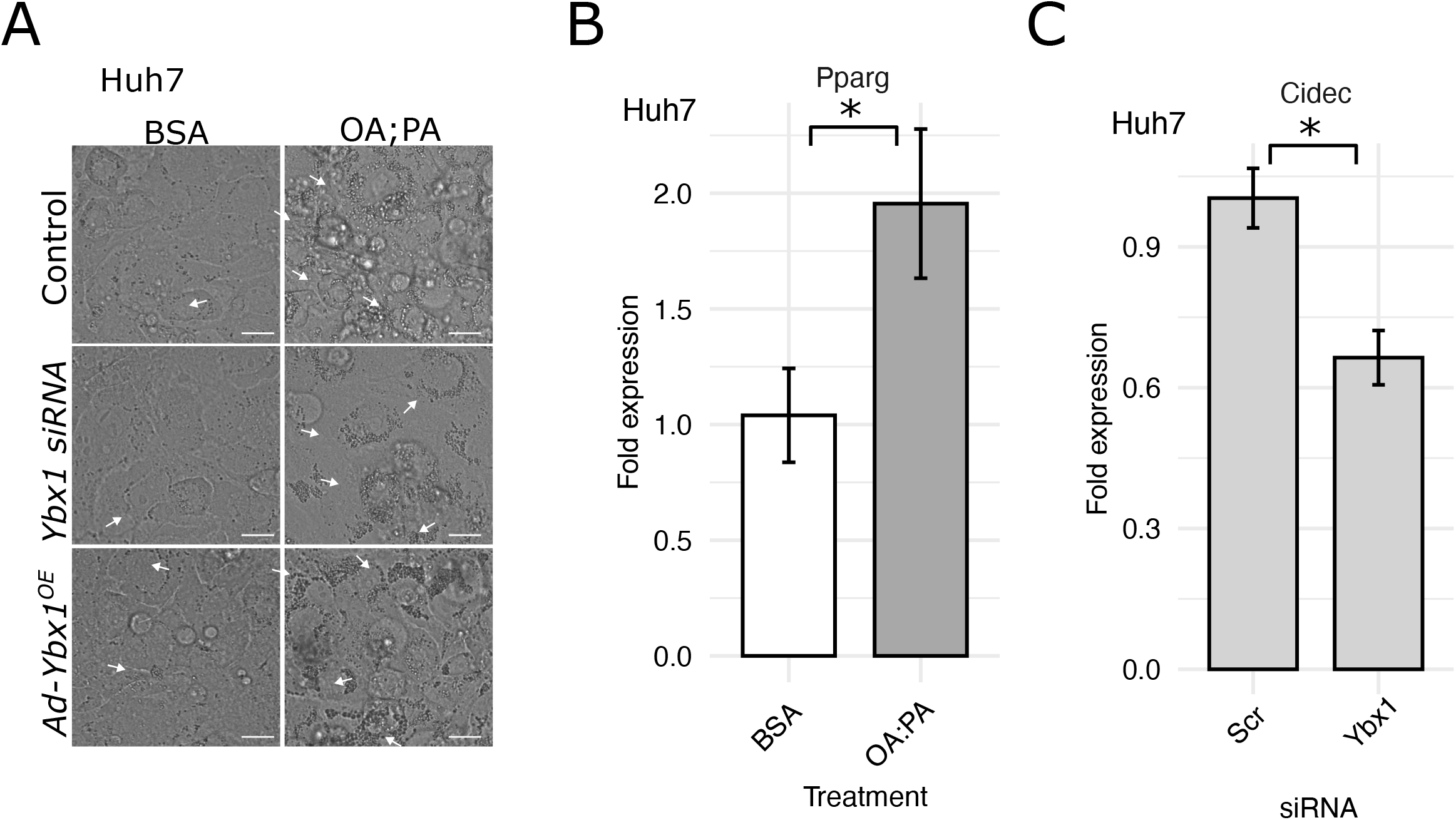

**Figure S4.**
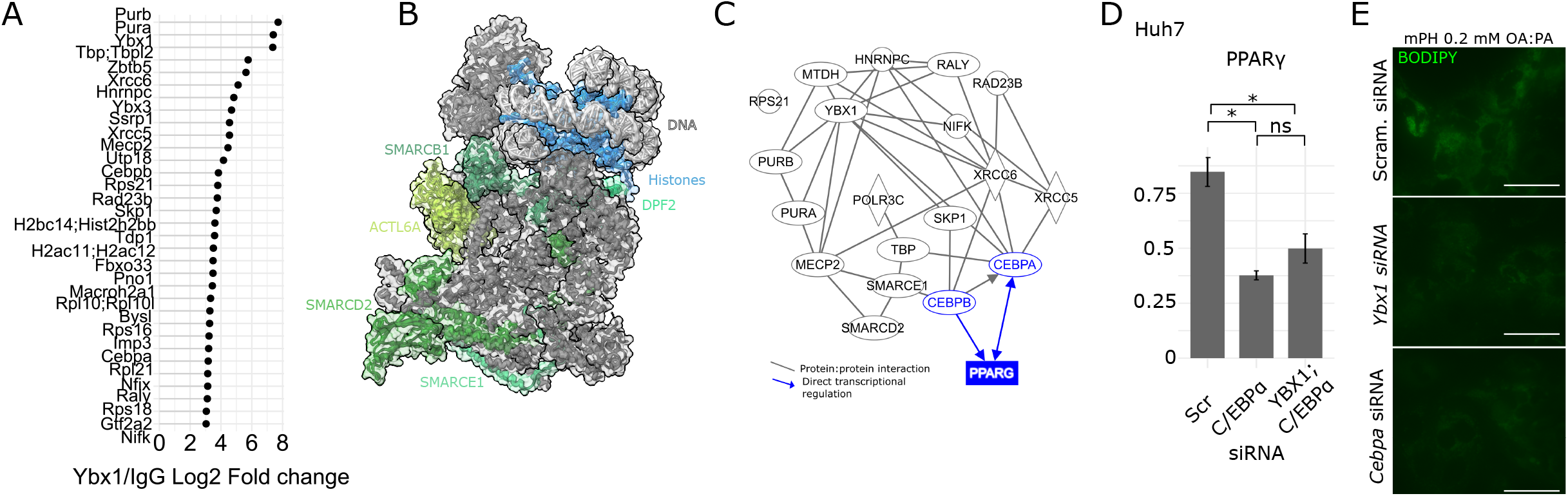

